# Genome-wide epigenetic profiling of 5-hydroxymethylcytosine by long-read optical mapping

**DOI:** 10.1101/260166

**Authors:** Tslil Gabrieli, Hila Sharim, Gil Nifker, Jonathan Jeffet, Tamar Shahal, Rani Arielly, Michal Levi-Sakin, Lily Hoch, Nissim Arbib, Yael Michaeli, Yuval Ebenstein

**Author notes:** These authors contributed equally to this work (Tslil Gabrieli, Hila Sharim).

## Abstract

The epigenetic mark 5-hydroxymethylcytosine (5-hmC) is a distinct product of active enzymatic demethylation that is linked to gene regulation, development and disease. Genome-wide 5-hmC profiles generated by short-read next-generation sequencing are limited in providing long-range epigenetic information relevant to highly variable genomic regions, such as the 3.7 Mbp disease-related Human Leukocyte Antigen (HLA) region. We present a long-read, single-molecule mapping technology that generates hybrid genetic/epigenetic profiles of native chromosomal DNA. The genome-wide distribution of 5- hmC in human peripheral blood cells correlates well with 5-hmC DNA immunoprecipitation (hMeDIP) sequencing. However, the long read length of 100 kbp-1Mbp produces 5-hmC profiles across variable genomic regions that failed to showup in the sequencing data. In addition, optical 5-hmC mapping shows strong correlation between the 5-hmC density in gene bodies and the corresponding level of gene expression. The single molecule concept provides information on the distribution and coexistence of 5-hmC signals at multiple genomic loci on the same genomic DNA molecule, revealing long-range correlations and cell-to-cell epigenetic variation.

## Introduction

The characterization and profiling of new DNA epigenetic modifications has been the focus of many studies in recent years. 5-hydroxymethylcytosine (5-hmC), the first in a chain of chemical oxidation products catalyzed by the Ten-eleven translocation (TET) family of dioxygenases during active DNA demethylation, has garnered special attention since its discovery in mammalian cells in 2009 ^12^. Although first thought to be a transient state, increasing evidence suggests that this modification plays a role in the regulation of gene expression, affecting development and cell differentiation ^3–7^. Additionally, depletion in global 5-hmC levels has been observed in various pathological conditions, including several types of cancer ^8–12^, making it a potential biomarker for early diagnosis and response to therapy.

The increasing interest in 5-hmC has served as a catalyst for the development of several techniques for profiling its distribution on a genome-wide scale. Currently accepted methods include single-base resolution methods such as oxidative bisulfite sequencing (oxBS-seq) ^13^ and TET-assisted bisulfite sequencing (TAB-seq) ^14^, as well as lower resolution affinity-based enrichment methods such as 5-hmC DNA immunoprecipitation (hMeDIP) ^4,15^ and 5-hmC selective chemical labeling (hMe-Seal) ^16^. The main limitation of all of these techniques is their reliance on next generation sequencing (NGS) which requires pooling and fragmentation of genomic DNA. As such, the epigenetic profile reported by these techniques is the average distribution of an entire population of cells, where cell-to-cell variation is lost, and with it the ability to detect small subpopulations. Recently reported single-cell 5-hmC sequencing may potentially capture 5-hmC variation^17^, but its low genomic sampling rate and reliance on NGS lead to difficulties in the analysis of long-range correlations at resonable cost^18^. Third generation sequencing approaches, including single-molecule, real-time (SMRT) sequencing (Pacific Biosciences), and nanopore sequencing (Oxford Nanopore Technologies) have demonstrated the potential ability to detect chemical modifications directly on single, long DNA molecules ^19–23^. However, extensive development of these applications is still necessary before they are used for whole-genome epigenetic profiling. A significant challenge arises when profiling tissues that exhibit ultra-low levels of 5-hmC such as human blood (0.001%- 0.005%). Specifically, gold-standard TAB-seq relies on enzymatic conversion that at best reaches 99% efficiency. In the case of blood, the result is an equivalent number of true and false 5-hmC signals, undermining the ability to extract informative data.

Here we present optical 5-hmC mapping, a single-molecule mapping approach for studying the genomic distribution of 5-hmC. We apply our method to human peripheral blood mononuclear cells (PBMCs), emphasizing the high sensitivity of this single molecule approach. Our method integrates into genome mapping technology, commercialized by BioNano Genomics Inc., which relies on extending fluorescently labeled DNA molecules in nanochannel arrays^24,25^. Fluorescence microscopy allows simultaneous detection of genetic and epigenetic information on the same molecule, by labeling each feature with a different color. Using color as a contrast mechanism, this method can be extended further to detect several epigenetic observables simultaneously, allowing the study of modification coexistence ^26,27^. In this study, a genetic barcode is generated by enzymatically labeling DNA in a specific sequence motif, and an additional epigenetic information layer is produced by labeling 5-hmC through a specific chemo-enzymatic reaction^28^. By aligning the genetic labels to a reference, the genomic positions of 5-hmC labels are mapped to obtain a genome-wide profile of epigenetic modifications (**Figure 1**).

**Figure 1.**
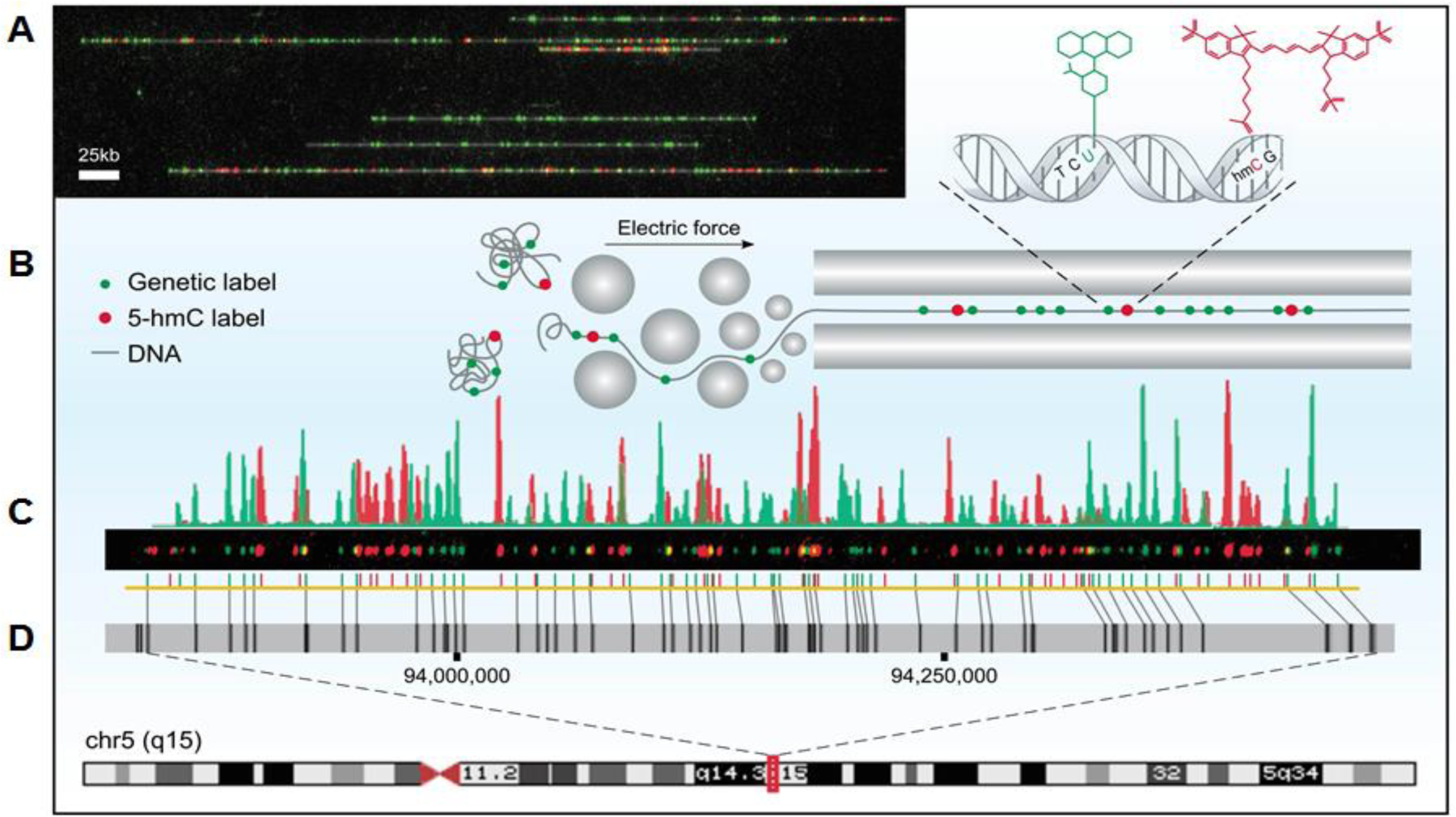
Optical 5-hmC mapping experimental scheme. **A.** Stretched DNA molecules (grey) fluorescently labeled in two colors. Green: sequence specific genetic barcode; Red: 5-hmC labels. **B.** Fluorescently labeled molecules are extended in nanochannel arrays by electrophoresis. **C.** Fluorescence intensity of genetic labels (green) and 5-hmC labels (red) along a single molecule. **D.** Genetic labels (green) are used to align a digital representation of the molecule in **C** (yellow) to an *in-silico* generated reference (grey) of chromosome 5, highlighting large structural variations such as the 7 kbp deletion in the mid-right part of the molecule, denoted by the diagonal alignment marks. 5-hmC labels (red) are mapped based on genetic alignment.

We compare the 5-hmC patterns detected by our optical mapping approach in PBMCs to hMeDIP-seq results generated for the same sample. We show that optical mapping displays higher sensitivity and lower background noise and that global patterns correlate well with previously reported results regarding the distribution of 5-hmC near regulatory elements and its enrichment in highly expressed genes. Furthermore, we show the correlation between 5- hmC density in gene bodies and their corresponding expression levels over a large dynamic range. Finally, we demonstrate the potential strength of long-read optical mapping in characterizing epigenetic patterns in variable genomic regions, which currently pose a challenge for NGS technology.

## Methods

### Human Subjects

The healthy donor sample used in this study was collected with informed consent for research use and approved by the Tel-Aviv University and Meir medical center ethical Review Boards, in accordance with the declaration of Helsinki.

### Measuring labeling efficiency of 5-hmC

For nicking reaction, 900 ng of lambda phage DNA (New England Biolabs) were digested with 30 units of Nt.BspQI nicking enzyme (New England Biolabs) for 2 hours at 50 °C in the presence of 3 μl 10X buffer 3.1 (New England Biolabs) and ultrapure water to a total volume of 30 μl. Next, nicked DNA was incubated for 1 hour at 72 °C with 15 units of Taq DNA polymerase (New England Biolabs), supplemented with 600 nM of dATP, dGTP, dCTP (Sigma) and atto-532-dUTP (Jena Bioscience), or dATP, dGTP, dTTP (Sigma) and 5hmdCTP (Zymo Research), in the presence of 4.5 μl 10X thermopol buffer (New England Biolabs) and ultrapure water to a total reaction volume of 45 μl. Following labeling, DNA was repaired for 30 minutes at 45 °C with 12 units of Taq DNA ligase (New England Biolabs) in the presence of 1.5 μl 10X thermopol buffer, 1 mM NAD+ (New England Biolabs) and ultrapure water to a total reaction volume of 60 μl. DNA that was labeled with 5hmdCTP nucleotides was further used for 5-hmC labeling. 300 ng of nicked DNA were incubated overnight at 37 °C with 30 units of T4 ß-glucosyltransferase (New England Biolabs), in the presence of 4.5 μl of 10X buffer 4 (New England Biolabs) and 200 mM homemade UDP-glucose-azide ^29^ in a total reaction volume of 45 μl. Next, Dibenzocyclooctyl (DBCO)-Cy5 (Jena Bioscience) was added to a final concentration of 620 μM and the reaction was incubated overnight at 37 °C. For purification of 5-hmC-labeled DNA from excess of fluorescent dyes, plugs were generated by adding agarose to the DNA (see “high molecular weight DNA extraction” section) to a final agarose concentration of 0.8%, and were then washed extensively with TE (pH 8) on a horizontal shaker. Plugs were melted and purified by drop dialysis using a 0.1 μm dialysis membrane (Millipore) floated on TE (pH 8). DNA from both labeling reactions was stained with YOYO-1 staining solution (BioNano Genomics) according to manufacturer’s instructions and gently mixed together with wide bore tips until homogenous. DNA concentration was measured by a Qubit HS dsDNA assay. Labeled DNA was loaded in nanochannels and imaged on an Irys system.

Images were processed by the IrysView software package (version 2.3, BioNano Genomics) to detect individual molecules and positions of labels along each molecule. In accordance with the length of the lambda phage genome (48.5 Kbp), only molecules that were 40 to 60 kbp in length were used for downstream analysis. Additionally, only labels with SNR ≥ 2.75 were considered. For each molecule, the number of 5-hmC labels (red channel) or the number of fluorescent nucleotides (green channel) were counted (in-house software). Molecules labeled in both colors were considered noise and discarded from the dataset.

### High-molecular-weight DNA extraction

Human peripheral blood mononuclear cells (PBMCs) were isolated from peripheral blood of a healthy donor by density gradient centrifugation using Ficoll Paque Plus (GE Healthcare) according to manufacturer’s instructions. PBMCs were trapped in agarose plugs to protect DNA from shearing during the labeling process ^30^. Samples were prepared according to the IrysPrep™ Plug Lysis Long DNA Isolation Protocol (Bionano Genomics Inc.) with slight modifications. Briefly, 1×10^6^ cells were washed twice with PBS, resuspended in cell suspension buffer (CHEF mammalian DNA extraction kit, Bio-Rad) and incubated at 43 °C for 10 minutes. 2% low melting agarose (CleanCut agarose, Bio-Rad) was melted at 70 °C followed by incubation at 43 °C for 10 minutes. Melted agarose was added to the resuspended cells at a final concentration of 0.7% and mixed gently. The mixture was immediately cast into a plug mold and plugs were incubated at 4 °C until solidified. Plugs were incubated twice (2 hours incubation followed by an overnight incubation) at 50 °C with freshly prepared 167 μl Proteinase K (Qiagen) in 2.5 ml lysis buffer (BioNano Genomics Inc.) with occasional shaking. Next, plugs were incubated with 50 μl RNase (Qiagen) in 2.5 ml TE (10 mM Tris, pH 8, 1 mM EDTA) for 1 hour at 37 °C with occasional shaking. Plugs were washed three times by adding 10 ml wash buffer (10 mM Tris, pH 8, 50 mM EDTA), manually shaking for 10 seconds and discarding the wash buffer before adding the next wash. Plugs were then washed four times by adding 10 ml wash buffer and shaking for 15 minutes on a horizontal platform mixer at 180 rpm at room temperature. Following washes, plugs were stored at 4 °C in wash buffer or used for labeling. In order to extract high molecular weight DNA, plugs were washed three times in TE pH 8, and were melted for 2 minutes at 70 °C, followed by 5 minutes incubation at 43 °C. Next, 0.4 units of Gelase (Epicenter) were added and the mixture was incubated for 45 minutes. High molecular weight DNA was purified by drop dialysis using a 0.1 μm dialysis membrane (Millipore) floated on TE (pH 8). Viscous DNA was gently pipetted and incubated at room temperature overnight in order to achieve homogeneity. DNA concentration was determined using Qubit BR dsDNS assay (Thermo Fisher Scientific).

### Genetic/epigenetic labeling and data collection

Genetic labeling was performed either using the commercial IrysPrep™ NLRS assay (Bionano Genomics Inc.) or an in-house developed alternative protocol (see “measuring labeling efficiency of 5-hmC” section). 900 ng of high molecular weight DNA from PBMCs was subjected to a nick-translation reaction with Nt.BspQI for nicking and atto-532-dUTP for labeling. For labeling of 5-hmC ^28^, nick-labeled DNA was incubated overnight at 37 °C with 90 units of T4 β-glucosyltransferase (New England Biolabs), in the presence of 13.5 μl 10X buffer 4 (New England Biolabs), 200 mM UDP-glucose-azide ^29^ and ultrapure water (135 μl total reaction volume). Next, Dibenzocyclooctyl (DBCO)-Cy5 (Jena Bioscience) was added to a final concentration of 620 μM and the reaction was incubated overnight at 37 °C. For purification of labeled DNA from excess fluorescent dye, plugs were prepared by adding agarose to the DNA as described above to a final agarose concentration of 0.8%. Plugs were then washed extensively with TE (pH 8) on a horizontal shaker. Plugs were melted and purified by drop dialysis as described, and DNA was stained with YOYO-1 staining solution (BioNano Genomics) according to manufacturer’s instructions with the addition of 25 mM Tris (pH 8) and 30 mM NaCl. DNA concentration was measured by the Qubit HS dsDNA assay (Thermo Fisher Scientific). For evaluation of photobleaching steps, labeled DNA was stretched on glass cover slips and imaged. For optical mapping experiments, labeled DNA was loaded into nanochannel-array Irys Chips and imaged on an Irys system (BioNano Genomics Inc.).

### Evaluation of 5-hmC sites per detected fluorescent spot

Due to the diffraction-limited resolution of optical mapping, a single fluorescent spot may contain more than a single 5-hmC site. We used single-molecule photobleaching to assess the distribution of sites per spot in the studied DNA sample, in order to correct for any bias caused by this potential underestimation. DNA molecules that were labeled with Cy5 to indicate 5- hmC locations and their backbone stained with YOYO-1 were stretched on modified coverslips prepared as previously described ^28^ with minor modifications. In short, 22 x 22 mm^2^ glass cover-slips were cleaned for 7 hours to overnight by incubation in a freshly made 2:1 (v/v) mixture of 70% nitric acid and 37% hydrochloric acid. After extensive washing with ultrapure water (18 MΩ) and then with ethanol, cover slips were dried under a stream of nitrogen. Dry cover-slips were immersed in a premixed solution containing 750 μl N-trimethoxysilylpropyl-N,N,N-trimethylammonium chloride and 200 μl of vinyltrimethoxysilane in 300 ml ultrapure water and incubated overnight at 65 °C. After incubation, cover-slips were thoroughly washed with ultrapure water and ethanol and stored at 4 °C in ethanol. The silane solution was freshly made and thoroughly mixed before the cover-slips were introduced into the mixture. Stored cover-slips were normally used within 2 weeks. Pre-labeled and stained DNA was diluted to a final concentration of 0.25 ng/μL in TE (pH 8) and 0.2 M DTT. DNA molecules were stretched on the silanized glass cover-slips by placing 6 μl of diluted DNA between a dry silanized coverslip and a non-treated microscope slide.

The extended DNA molecules were imaged on an Olympus IX81 microscope adapted for laser-fluorescence microscopy. A 637 nm CW laser diode (Coherent, OBIS 637LX) and a 473 nm CW laser (OEM, 200mW) were used as excitation sources for the 5-hmC Cy5 labels and YOYO-1 intercalating backbone staining dye, respectively. Excitation light was focused on the back aperture of a 100x oil immersion objective (Olympus, UPlanSApo 100x/N.A 1.4) with an average excitation power density of 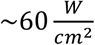 in the sample plain. Sample fluorescence was collected by the same objective and imaged through a polychroic beamsplitter (Chroma, ZT473/532/637 rpc-xt890) and emission filter (Semrock, 679/41 Brightline) onto an electron multiplying charge coupled device (EMCCD) camera (Andor, iXon Ultra 897) with singlemolecule detection sensitivity. All measurements were carried out at room temperature under ambient conditions. Upon excitation, 5-hmC sites were detected as fluorescent diffraction limited spots in the field of view (80 × 80 *μm*^2^). Time lapse movies (500 frames, 20 ms/frame) were acquired for over 100 different fields of view, containing 1000 individual isolated observable 5-hmC markers sites. For each time lapse movie, individual labels were automatically localized by 2D Gaussian fitting and manually validated to be isolated fluorescent marks consisting of a single Gaussian spot and co-localized with a DNA molecule. After localization, time traces of fluorescence intensity were calculated for each marker. The intensity was calculated by the average intensity of a 3×3 pixel^2^ window around the center of each fitted Gaussian. In order to account for the uneven excitation intensity a local background subtraction was performed for each marker by taking the average intensity of a 7×7 pixel^2^ contour window around the fitted Gaussian.

The number of photobleaching steps was manually quantified according to the number of intensity drops in each time-trace. All data analysis in this section was carried out by a custom made MATLAB program.

### Optical mapping and data analysis

Two optical epigenome mapping experiments of PBMCs were performed separately, and their results were combined for downstream analysis.

Raw images were processed by the IrysView software package (version 2.3, BioNano Genomics Inc.) to detect individual molecules and positions of genetic and epigenetic labels along each molecule. Only labels with SNR ≥ 2.75 were considered. The positions of genetic labels were used to align molecules longer than 150 kbp to an *in-silico* generated map of expected Nt.BspQI nicking sites in the hg19 human reference (IrysView), and only molecules with an alignment confidence equal or higher than 12 (P <= 10^−12^) were used for downstream analysis. In order to minimize ambiguous alignments of the overlaid 5-hmC pattern, we filtered out molecules that were not aligned to the reference for over 60% of their total length. Genomic positions of epigenetic labels were then extracted by interpolation, based on the genetic alignment results (in-house script). 500 bp were added to either side of each label position, in order to account for optical resolution, and genomic coverage of both 5-hmC and aligned molecules was calculated using BEDTools (version 2.25.0)^31^ genomecov. Regions represented by less than 19 molecules were discarded for downstream analysis and molecule coverage was normalized by dividing the coverage in each genomic position by the maximum coverage. Normalized 5-hmC coverage was then calculated by dividing the number of detected 5-hmC labels in each genomic position by the normalized molecule coverage in the same position.

### hMeDIP-seq library preparation

DNA extracted from PBMCs was used to prepare two sequencing libraries by Epigentek Inc. Each library included one hMeDIP sample and an input sample. hMeDIP experiments were performed with 5-hmC monoclonal antibody, using 1.5-4 μg DNA. Sample libraries were sequenced on a HiSeq 2500 (Illumina Inc.).

### Sequencing alignment and data analysis

Sequencing reads were aligned to the hg19 human reference using Bowtie 2 (version 2.2.6) ^32^ with default parameters. Following alignment, reads with MAPQ less than 30 were filtered with SAMtools^33^, and the remaining reads were de-duplicated with Picard (https://broadinstitute.github.io/picard/) to eliminate PCR duplication bias. Reads from the pull-down data set were extended in the 3’ direction to a total length of 200 bp, in order to account for insert size. Genomic coverage of 5-hmC and input DNA was calculated using BEDTools genomecov. Due to a low signal to noise ratio in the pull-down experiment, the signal extraction scaling (SES) ^34^ method was used to estimate a minimal signal level. Briefly, the genome was divided into 1 kbp non-overlapping windows and the number of 5-hmC and input reads in each window was counted. Windows were then sorted in increasing order based on 5-hmC counts, and the cumulative sum in each window was calculated and normalized to the cumulative sum of the complete data set. The absolute value of the difference between these normalized sums was calculated for each window, and the minimal signal level was defined as the 5-hmC counts value in the window where this difference reached a maximum. Based on this calculation, all regions where the 5-hmC signal was lower than 7 were set to zero. Additionally, ENCODE-defined signal artifact regions in the hg19 human genome reference ^35^ (https://sites.google.com/site/anshulkundaje/projects/blacklists) were discarded.

For visual assessment of the correlation between the hMeDIP-seq results and the optical mapping results, sequencing coverage resolution was lowered to 1 kbp using a running average calculation. This was accomplished by dividing the genome into overlapping 1 kbp windows with a 1 bp shift and calculating the mean 5-hmC coverage in each window. The calculated mean was then set as the 5-hmC value in the midpoint genomic position of each window.

### Analysis of global 5-hmC distribution across genomic features

Transcription start sites (TSS) were defined according to RefSeq annotation for hg19 downloaded from the UCSC genome browser database. H3K4me1 and H3K27Ac ChIP-seq data for PBMCs was downloaded as alignment results from the Roadmap Epigenomics Project (GEO accession numbers GSM1127143 and GSM1127145, respectively), and converted into genomic coverage using BEDTools. Enhancer regions were defined by examining coverage percentiles (>99% percentile for H3K4me1 and >99.5% for H3K27Ac), and regions closer than 100 bp apart were merged. 5-hmC counts in each region were calculated using BEDTools by dividing regions into equally sized bins (90 bp for H3K4me1, 30 bp for H3K27Ac, and 20 bp for TSS), computing the mean 5-hmC signal in each bin, and summing the signal across all bins of equal distance from the region midpoint. Normalization of values to a 0-1 range was performed separately for each data set.

### Analysis of 5-hmC correlation to gene expression

Expression data of protein-coding genes for PBMCs was downloaded from the Roadmap Epigenomics Project as RPKM values

(http://egg2.wustl.edu/roadmap/data/byDataType/rna/expression/). Genes were classified into 3 groups: high expression (5987 genes, log_*10*_ *RPKM) ≥* 10), low expression (5741 genes, 1 ≤ log _*10*_*(RPKM) <* 10), and no expression (8074 genes, log_*W*_ *(RPKM)* < 1). Mean 5-hmC values along genes were calculated using deepTools^36^ computeMatrix in scale regions mode. Each gene was scaled to 15 kbp and divided into 300 bp bins. The mean 5-hmC score in each bin was calculated, and all scores in the same bin were summed and normalized to the number of genes in the group.

## Results

We developed optical 5-hmC mapping on high molecular weight DNA extracted from fresh human PBMCs. A nick-translation reaction with the nicking enzyme Nt.BspQI was performed in order to incorporate a green fluorophore into the DNA, producing a sequence specific labeling pattern for genome mapping. A second layer of information was obtained by performing a chemo-enzymatic reaction to specifically label 5-hmC with a red fluorophore.

### Efficiency of 5-hmC labeling

Efficient labeling of 5-hmC is critical for obtaining meaningful information from individual molecules. In order to evaluate the efficiency of 5-hmC labeling, we performed two separate nick-labeling reactions on purified lambda phage DNA. The reactions contained either fluorescent nucleotides, for assessment of nick-labeling efficiency, or 5-hmC nucleotides, which were then fluorescently labeled according to our 5-hmC labeling scheme and used for assessment of 5-hmC labeling efficiency. DNA from both reactions was combined, driven into nanochannels, and imaged together on an Irys instrument (Figure 2A). The Lambda phage genome contains 10 expected Nt.BspQI nicking sites, with two sites that cannot be separated due to the optical resolution limit. By comparing the detected spots to the expected nicking sites, labeling efficiency was calculated.

**Figure 2.**
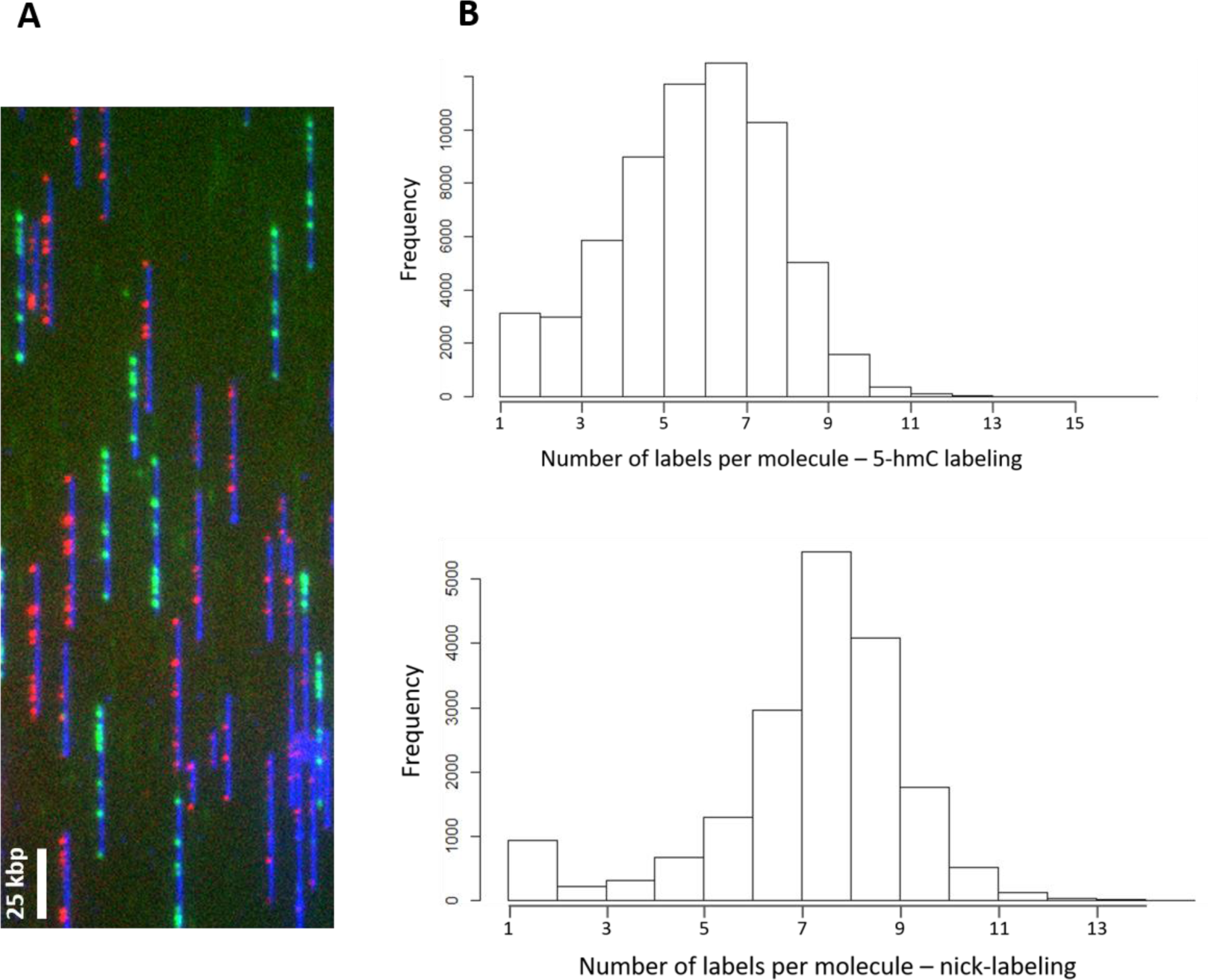
Assessment of 5-hmC labeling efficiency. Lambda DNA was nicked with Nt.BspQI (10 expected nicking sites) and labeled with either 5-hmC or fluorescent dUTP. 5-hmC was labeled according to our labeling scheme and the samples were mixed and imaged together in order to evaluate the labeling efficiency. **A.** representative field of view showing a mixed population of green (nicking) and red (5- hmC) labeled molecules. **B.** Histograms showing the number of labels per molecule for 5-hmC labeling (top) and nicking (bottom).

Figure 2B shows the number of dots per molecules in the red channel (5-hmC labeling) and in the green channel (nick-labeling). Nick-labeling efficiency was calculated by dividing the average number of green labels per molecule in each scan by the total number of expected labels. 5-hmC labeling efficiency was calculated by dividing the average number of red labels per molecule by the average number of green labels per molecule in each scan. 20,520 scanned images were analyzed to determine labeling efficiency. Accordingly, the nicking efficiency was determined as 80-88%, and the 5-hmC labeling efficiency as 78-87%.

### Quantification of 5-hmC sites per detected label by measuring photobleaching steps

The number of fluorophores detected in each isolated 5-hmC label has a crucial role in determining the actual density of 5-hmC. A scenario in which an isolated label is comprised of many fluorophores, due to multiple 5-hmC sites residing in a sub-resolution area along the DNA, could have a significant impact on the validity of our analysis. Therefore, it is crucial that we validate the characteristic amount of fluorophores, and therefore the number of 5-hmC sites, per isolated fluorescent label. In order to assess the number of fluorophores, we monitored the photobleaching process which fluorophores undergo when exposed to intense laser excitation (see methods). Quantifying the amount of photobleaching steps enabled us to accurately determine the number of 5-hmC sites per detected spot (Figure S1-S3). Approximately 1000 fluorescent spots along the DNA were analysed in order to construct the distribution of 5-hmC clusters. The full distribution allowed us to correct for this effect as part of the calibration procedure. Effectively, 1.45 fluorophores are present in each labeling site and this correction factor was taken into account for global quantification. Since Over 90% of the analyzed 5-hmC spots contained 1-2 5-hmC labels, our method can reliably quantify 5- hmC in genomic DNA from blood, without considering the intensity of the labels. Using such global quatification we have recently shown that our labeling scheme can detect a decrease in the global level of 5-hmC in blood and colon tumor cells vs. normal cells^12^.

### Generation of genome-wide 5-hmC profiles

To recover the genome-wide distribution of 5-hmC we combine our labeling scheme with conventional genome mapping by generating a sequence specific fluorescent barcode in addition to the epigenetic marks. This dual-clolr labeling provides information on the amount, as well as the location and distribution of 5-hmC sites along the genome.

Labeled DNA was stretched and imaged on nanochannel array chips, and the positions of genetic and epigenetic labels were automatically detected. Molecule images and genome browser compatible molecule tracks were generated with IrysExtract ^37^ (Figure 1A, 1C). 992030 long molecules (>150 kb) were aligned to the reference (human hg19), and 587583 that passed our confidence criteria (see methods) were used for downstream analysis, generating a median genomic coverage of 46X. In order to verify the 5-hmC density obtained from optical mapping, we performed LC-MS/MS measurements on DNA extracted from the same PBMCs sample (see supporting information, Figure S4). The percentage of 5-hmC in DNA extracted from PBMCs was calculated as 0.0035%. This is in agreement with the mean 5- hmC density measured for the optical mapping experiment (0.0029%).

In order to validate our optical mapping results, we also performed two independent hMeDIP-seq experiments on the same sample. Based on the correlation between the genomic 5-hmC profiles of these two experiments (Pearson correlation coefficient = 0.7) we were able to merge the sequencing reads from both datasets, in order to enhance the 5-hmC signal. The extremely low levels of 5-hmC in blood ^38^, together with non-specific pulldown inherent in this type of experiments ^39,40^, resulted in low signal-to-noise ratio for the sequencing data. Thus, a minimal threshold for 5-hmC reads was set for downstream analysis (supporting information, Figures S5-S6).

### Global optical mapping patterns of 5-hmC correlate with sequencing results

We first wanted to verify that the epigenetic patterns we observe using our optical mapping approach correlate with the patterns observed in the sequencing results. To this end, we examined the global distribution of 5-hmC in both data sets near several regulatory elements (Figure 3A-C). Both methods display distinct patterns near transcription start sites (TSS), enhancers marked by the histone modification H3K4me1, and active enhancers marked by the histone modification H3K27Ac, as previously reported for embryonic stem cells (ESCs), brain, and blood ^41–43^. In all cases, the patterns detected by both methods are highly correlated. However, global examination of both datasets on a genome-wide scale (Figure 3) establishes the advantages of the optical mapping approach. While the sequencing results show finer features due to the higher resolution of hMeDIP (~200 bp) compared to optical mapping (~1500 bp), the optical data displays superior signal-to-noise ratio allowing us to reliably detect low levels of 5-hmC. This high sensitivity is likely due to the low false positive and false negative rates of our labeling scheme (Figure 1). Moreover, while antibody-based methods are biased towards heavily modified regions ^40^, the high sensitivity of optical detection, combined with the single-molecule aspect of this approach, allow the detection of rare, isolated 5-hmC residues occurring only in a small subset of cells. This is clearly seen in the additional signals present in the optical mapping track in figure 3D.

**Figure 3.**
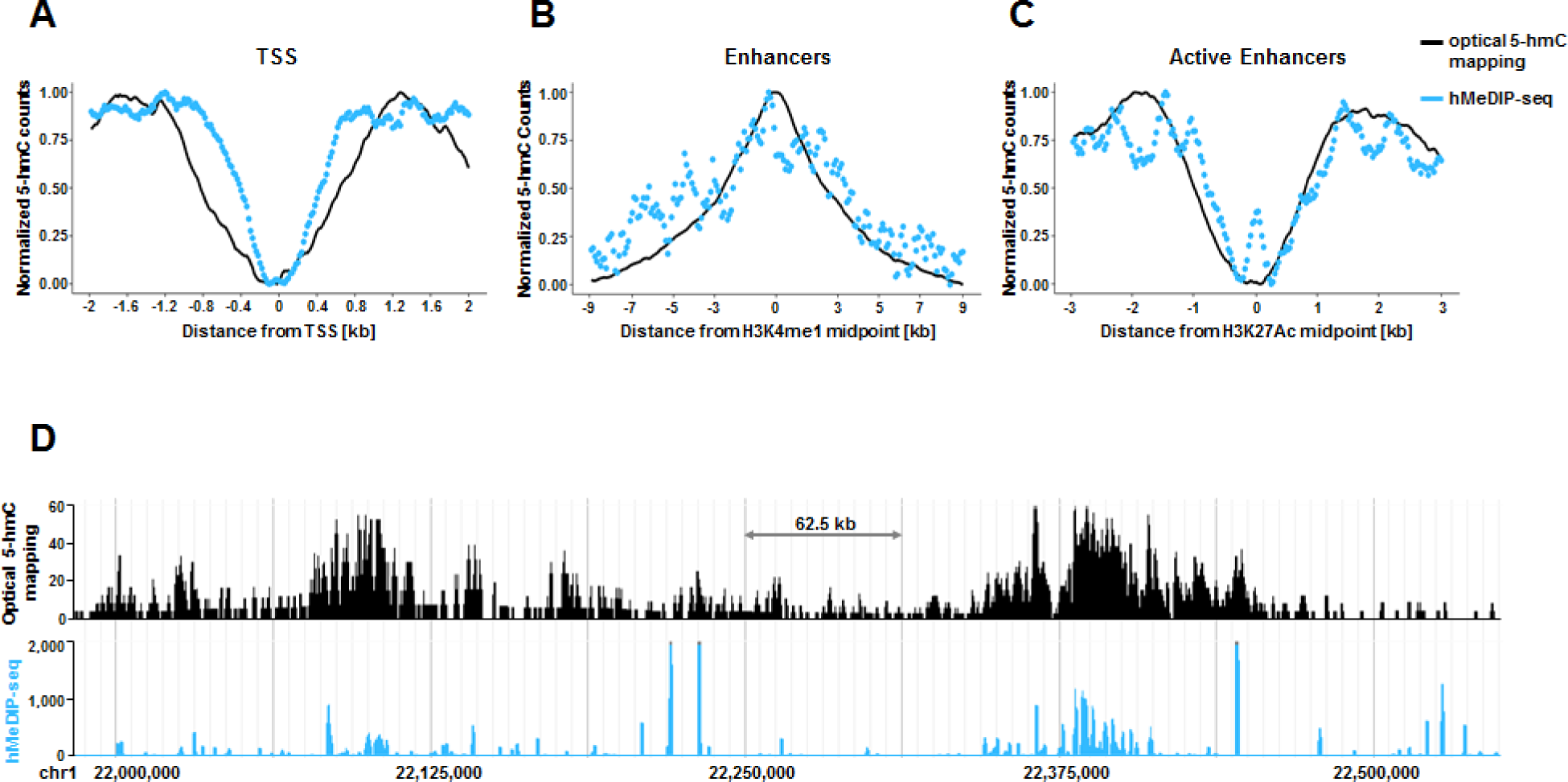
Global correlation between 5-hmC profiles produced by optical 5-hmC mapping and hMeDIP- s eq. **A.** Coverage as a function of distance from TSS. **B.** Coverage as a function of distance from H3K4me1 histone modification peaks. **C.** Coverage as a function of distance from H3K27Ac histone modification peaks. **D.** Comparison of coverage produced by both methods in a representative 500 kbp region from chromosome 1. hMeDIP-seq results are presented in 1 kbp resolution.

### 5-hmC levels in gene bodies correlate with gene expression

Following reports regarding 5-hmC enrichment in the bodies of highly expressed genes ^41–43^, we examined the correlation between gene expression and 5-hmC levels for both optical mapping and hMeDIP-seq results (Figure 4). Although for both methods higher levels of 5- hmC are observed in highly expressed genes, optical 5-hmC mapping presents a much larger dynamic range. The high sensitivity and specificity of the fluorescent labeling utilized in optical 5-hmC mapping enables clear distinction between the 5-hmC levels in gene bodies of low expressed and unexpressed genes. These two groups are almost indistinguishable in the hMeDIP-seq results. These results suggest that 5-hmC levels may be used to infer gene expression without directly quantifying RNA levels.

**Figure 4.**
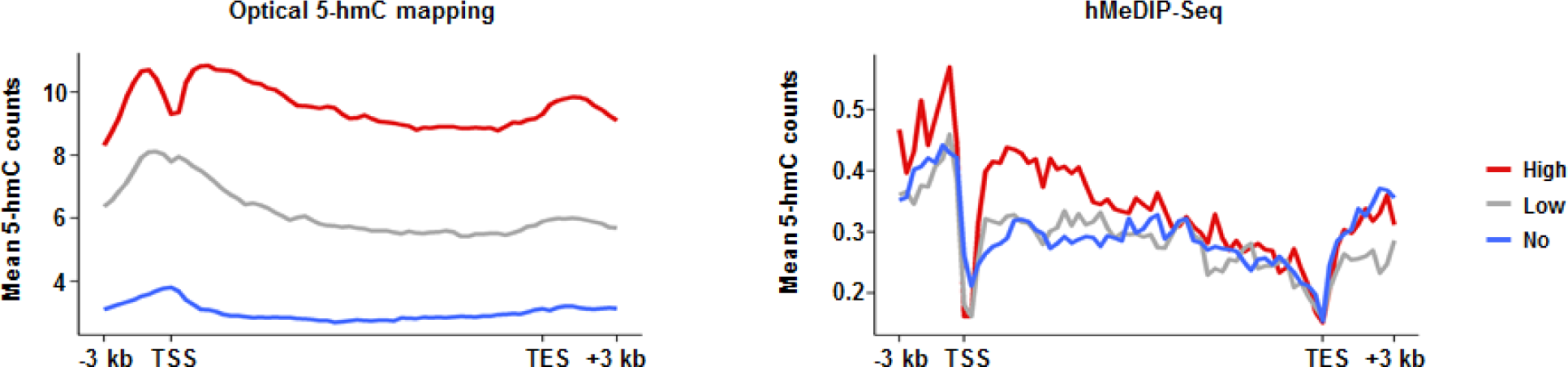
5-hmC coverage across gene bodies, in correlation with gene expression level. Left: optical mapping data. Right: hMeDIP-seq data.

### Optical mapping “long reads” allow the characterization of highly variable regions

One of the main advantages of genetic optical mapping is the ability to use long-range information encoded in long DNA molecules to characterize highly variable genomic regions ^44,45^. These long-reads extend beyond the variable region and may be anchored to the reference based on reliable alignment to conserved flanking regions, maintaining contiguity along the variable region itself.

The Human leukocyte antigen (HLA) region, located on chromosome arm 6p21.3, is one of the most polymorphic regions in the human genome ^46^. This 3.6 Mb region has been associated with more than 100 diseases, including diabetes, psoriasis, and asthma, and several alleles of HLA genes have been linked to hypersensitivity to specific drugs ^47,48^. Therefore, information regarding the epigenetic landscape of this region may have clinical importance.

Examining the region around the HLA-A gene, one of three major histocompatibility complex (MHC) class I cell surface receptors, we found that single molecules can indeed be aligned to the reference with high confidence around this locus (Figure 5A). This resulted in the detection of high levels of 5-hmC in this region, information that was not detected by hMeDIP-seq as there were no sequencing reads aligned to this gene (Figure 5B). The highly variable nature of this locus is known to pose a challenge to NGS sequencing technologies. In contrast, the alignment of long optical reads to the reference is not hampered by the variable region since it occupies only a small portion of the detected molecule. Consequently, using optical 5-hmC mapping we are able to directly detect 5-hmC modifications in the HLA region without the amplification or specific targeting methods currently needed for short-read-based experiments ^49,50^.

**Figure 5.**
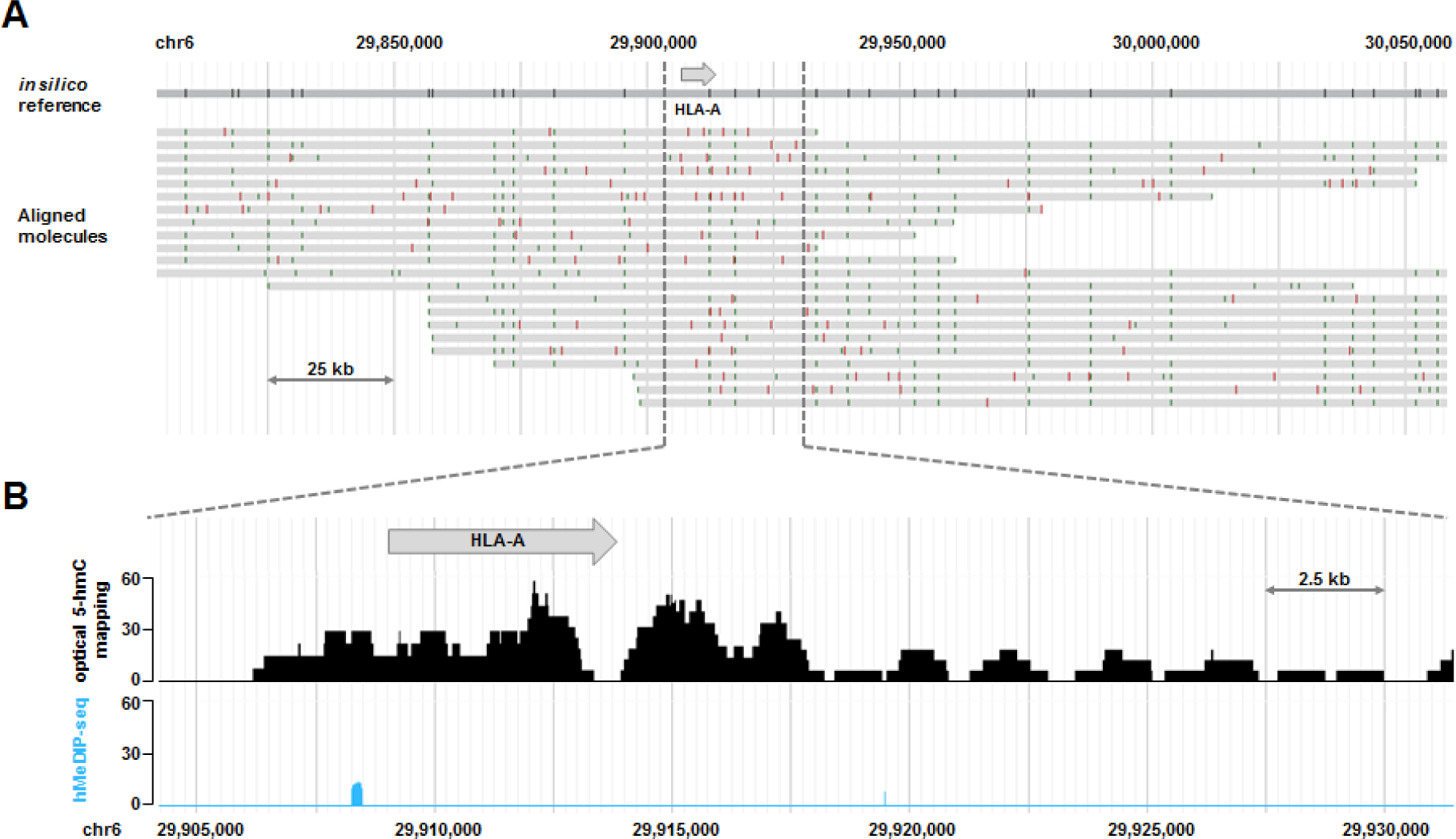
Epigenetic characterization of the variable HLA region by optical 5-hmC mapping “long reads”. **A.** Single molecules (grey) aligned to an *in silico* reference in a 300 kbp segment of the HLA region. Green: sequence specific genetic barcode; Red: 5-hmC labels. **B.** 5-hmC coverage of 25 kbp around the HLA-A gene. Black: optical 5-hmC mapping; Blue: hMeDIP-seq.

## Discussion

In recent years, genetic optical mapping has proven to be invaluable for the characterization of variable and repetitive regions inaccessible to NGS technologies, as well as for completing assemblies of various species ^51,52^. In this work we demonstrate the ability to add a second, epigenetic layer of information to long individual DNA molecules. The combination of existing optical genome mapping technology with a 5-hmC-specific labeling reaction creates a genome-wide profile of this epigenetic mark in human PBMCs.

We demonstrate that optical 5-hmC mapping produces an epigenetic map that correlates well with the epigenetic map produced by hMeDIP-seq, which relies on gold-standard NGS technologies. Despite the resolution limit of optical detection, optical 5-hmC mapping proves to be superior to hMeDIP-seq in sensitivity and specificity, resulting in a high signal-to-noise ratio and broad dynamic range. Our labeling and imaging technique highlights single 5-hmC sites with no false positives and a minor false negative rate. These attributes enable the detection of extremely low levels of epigenetic modifications. This is especially important for cancer related studies, where 5-hmC levels decline drastically with disease progression ^8,9,11,53^ Additionally, utilizing the long range information encoded in optical “long reads” enables the epigenetic characterization of variable and repetitive regions known to pose a challenge to NGS technologies. The single-molecule aspect of optical 5-hmC mapping preserves the ability to characterize single cells by examining molecules originating from the same genomic loci. Although single-cell 5-hmC sequencing provides information on 5-hmC distribution in single cells, it samples only ~10% of 5-hmC per cell. This low sampling efficiency prevents this technique from successfully characterizing the 5-hmC distribution of selected genomic loci without sequencing thousands of single-cells. Rather, it is useful mainly for analysis on a chromosome scale ^17^. TAB-seq, another gold standard technique for genome wide 5-hmC profiling, can detect the epigenetic mark with high resolution and high efficiency ^14^. Nevertheless, it is limited to tissues expressing relatively high levels of 5-hmC. This is due to the conversion rate of 5-methylcytosine (5-mC) to 5-carboxycytosine (5-CaC) by the TET enzyme. Even in the case of 99% conversion efficiency, 1% of non-converted methylated C will be detected as 5-hmC. This false positive rate of TAB-seq is in the same order of the real 5- hmC content in tissues exhibiting extremely low levels of 5-hmC such as blood. With an average signal to noise ratio of 1, TAB-seq is not suitable for characterizing the distribution of 5-hmC in PBMCs.

Finally, it is important to note that the presented concept is not limited to the measurement of 5-hmC, or to the measurement of a single epigenetic feature. As fluorescence microscopy relies on color to distinguish different genomic features, the number of markers that can be detected simultaneously is limited only by the number of specific labeling chemistries and the spectral properties of the optical measurement system. Thus, further development of this technique may enable the simultaneous measurement of 5-hmC and 5-mC, giving a more comprehensive genomic profile of the cell population as a whole, and of cell-to-cell variability in particular.

